# Human cell surface-AAV interactomes identify LRP6 as blood-brain-barrier transcytosis receptor and immune cytokine IL3 as AAV9 binder

**DOI:** 10.1101/2024.01.05.574399

**Authors:** Timothy F. Shay, Seongmin Jang, Xinhong Chen, Beth Walker, Claire Tebbutt, Damien A. Wolfe, Tyler J. Brittain, Cynthia M. Arokiaraj, Erin E. Sullivan, Xiaozhe Ding, Ting-Yu Wang, Yaping Lei, Miguel R. Chuapoco, Tsui-Fen Chou, Viviana Gradinaru

## Abstract

Adeno-associated viruses (AAVs) are foundational gene delivery tools for basic science and clinical therapeutics. However, lack of mechanistic insight, especially for engineered vectors created by directed evolution, can hamper their application. Here, we adapted an unbiased human cell microarray platform to determine the extracellular and cell surface interactomes of natural and engineered AAVs. We identified a naturally-evolved and serotype-specific interaction between the AAV9 capsid and human interleukin 3 (IL3), with possible roles in host immune modulation, as well as lab-evolved low-density-lipoprotein-receptor-related-protein 6 (LRP6) interactions specific to engineered capsids that cross the blood-brain barrier in non-human primates after intravenous administration. The unbiased cell microarray screening approach also allowed us to identify off-target tissue binding interactions of engineered brain-enriched AAV capsids that may inform vectors’ peripheral organ tropism and side effects. These results allow confident application of engineered AAVs in diverse organisms and unlock future target-informed engineering of improved viral and non-viral vectors for non-invasive therapeutic delivery to the brain.

## Introduction

Adeno-associated viruses (AAVs) have become the gene delivery vector of choice at the bench and in the clinic^1,2^. Systemic administration of AAVs, such as AAV9^3–6^, allows noninvasive gene delivery, particularly in large or distributed biological structures^7^, but access to the brain from the periphery is restricted by the blood-brain barrier (BBB), a complex biological structure that regulates molecular access to the central nervous system (CNS)^8–10^. Systemic administration of AAVs also exposes the vectors to the host immune system^11,12^ and off-target tissues^3,13^. The poor efficiency of brain targeting after systemic administration with natural serotypes often necessitates high doses that raise costs and may trigger serious adverse events^14–16^. Thus, improved vectors are needed if AAV gene therapy is to realize its full therapeutic potential.

AAV capsid engineering, particularly through directed evolution methods, has demonstrated that markedly improved efficiency in desired cell types and tissues after systemic intravenous delivery is possible^17–19^. In particular, two recently identified engineered capsids, AAV9-X1.1^20^ and AAV.CAP-Mac^21^, robustly transduce CNS neurons after systemic administration in macaque. As AAV capsids are applied across species however, the enhanced tropisms of many engineered vectors can vary^22,23,20,21^. This is concerning for human clinical trials, as a capsid developed in non-human species that performs poorly when translated to humans may not only fail to provide therapeutic benefit but might preclude that patient from future therapies by inducing neutralizing antibodies^11^.

This translational challenge of AAV engineering through directed evolution also represents an opportunity to better understand fundamental mechanisms of drug delivery to the brain. Directed evolution of engineered capsids with enhanced BBB crossing provides a platform with which researchers may survey the most efficient pathways across this barrier. While recent progress suggests that engineered AAVs may utilize diverse BBB-crossing receptors^24–27^, the mechanisms of primate brain-enhanced vectors^20,21,23,28–30^ remain underexplored.

To address this challenge, we adapted Retrogenix cell microarrays^31,32^ of the human membrane proteome and secretome to screen natural and engineered AAV capsid interactions with host cells. This allowed us to rapidly assay more than 90% of the human membrane proteome and secretome, including key protein classes such as receptors, transporters, and cytokines. Using this broad, unbiased screen, we identified several new AAV interactions with implications for the host immune response (human interleukin 3 (IL3) binding to AAV9), enhanced BBB crossing across species (via low-density-lipoprotein-receptor-related-protein 6 (LRP6) binding by AAV9-X1.1 and CAP-Mac), and peripheral tissue tropism (through pancreas-expressed glycoprotein 2 (GP2) binding by AAV9-X1.1 and CAP-Mac). Understanding the mechanism of action of systemic AAVs through methods such as those used here will be critical for successful vector translation and enables design of improved vectors, as well as other therapeutic protein modalities, for specific targets^33,34^.

## Results

### High-throughput screening for AAV binding partners

To screen AAV-binding proteins, we used Retrogenix cell microarrays^31,32^ of the human membrane proteome and secretome, in which DNA oligos encoding human membrane and secreted proteins are affixed at known slide locations (Figure 1a). HEK293 cells are then grown on the slides and become individually reverse-transfected with the oligos in the corresponding pattern. AAVs that directly interact with a given protein will preferentially bind to cells expressing that protein; other slide locations define non-specific background binding. To increase confidence in binding specificity, each protein is patterned at two different locations (four locations presented for initial condition optimization) (Figure 1b and c). We optimized screen conditions using previously-identified AAV and interacting protein pairs, (1) AAV9 with AAVR (KIAA0319L)^35^ and (2) PHP.eB with mouse LY6A^25–27^, for two different detection methods: biotin tagging and antibody direct detection (Figure 1b and 1c). Biotinylated capsids can be detected with fluorescent streptavidin, and unlabeled capsids are detected with an antibody whose epitope is distinct from the commonly engineered capsid variable regions IV and VIII^18,36^. As noted previously^37,38^, capsid primary amine labeling levels must be tuned so that surface modification does not interfere with capsid key binding interactions. We found that the best signal to noise ratio for duplicate spots (calculated as the average intensity across positive control spots compared to the average intensity of the rest of the slide) was achieved by directly fixing cell-bound AAVs without washes.

**Figure 1.**
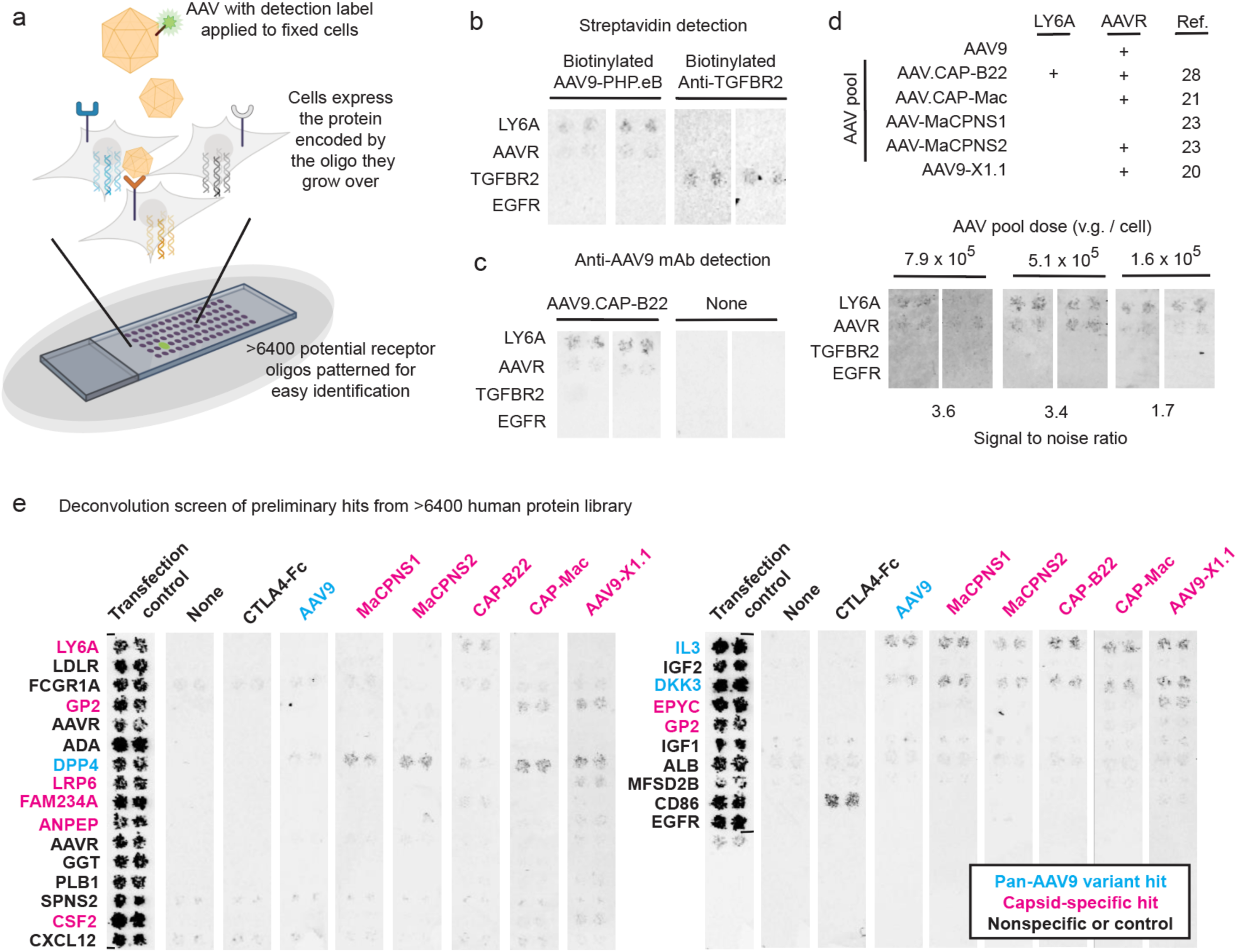
High-throughput screen identifies AAV-binding human proteins. ***a***, Schematic of AAV cell microarray screen. DNA oligos that encode individual membrane proteins are chemically coupled to slides in a known pattern, reverse transfecting the cells that grow on them and thereby creating spots of cells overexpressing a particular, known protein. Each protein is expressed in duplicate at two different slide locations on the slide. When AAVs are applied to the slides, enhanced binding can be detected from duplicate cell spots overexpressing cognate AAV receptors. ***b***, Known AAV capsid receptor interactions, such as AAVR and LY6A with AAV-PHP.eB, were used to optimize conditions for streptavidin-based detection of biotinylated capsids with two sets of replicate spots. Anti-TGFBR2 antibody was used as a non-AAV positive control. ***c***, AAVR and LY6A interaction with AAV9.CAP-B22 were used to optimize conditions for anti-AAV9 antibody direct detection of unmodified capsids with two sets of replicate spots. Anti-TGFBR2 antibody was used as a non-AAV control. ***d***, Pooled AAV capsid screening conditions were optimized by varying the concentrations of individual capsids within the pool to maximize signal to noise after direct detection with anti-AAV9 antibody, with two sets of replicate spots. ***e***, Pooled screening led to preliminary hits which were deconvoluted by individual-capsid screens, identifying novel potential capsid-binding proteins by direct detection with anti-AAV9 antibody. Transfection control condition detected fluorescent protein reverse transfected along with each receptor. None condition was treated only with anti-AAV9 antibody. Proteins in cyan were identified in all individual AAV screens, and likely represent interactions outside the engineered regions of AAV9. Proteins in magenta specifically bind to at least one engineered capsid.

We proceeded with antibody direct capsid detection and validated conditions with a panel of AAV capsids, including AAV9 as well as five engineered AAV9 variants with enhanced potency in the CNS of non-human primates (NHPs) after systemic administration (AAV.CAP-B22^28^, AAV.CAP-Mac^21^, AAV-MaCPNS1^23^, AAV-MaCPNS2^23^, and AAV9-X1.1^20^)(Table 1 & Figure 1d)^20,21,23,28^. Testing these capsids individually revealed that all AAVs except MaCPNS1 exhibited detectable AAVR binding (the exception may be due to the interfering geometry of the capsid’s variable region VIII insertion^39^), whereas only CAP-B22 interacted with mouse LY6A (likely through the PHP.eB loop in variable region VIII^25–27)^ (Figure 1d and Extended Data Figure 1). To enable higher-throughput screening, we decided to test the six capsids as a pool. Pooled testing required additional dosage optimizing, first for the individual and then for the collective background binding levels of the included capsids (Extended Data Table 1). An optimal dose was determined that minimized background binding while still allowing the unique interaction of CAP-B22 with mouse LY6A to be distinguished from the five non-LY6A-interacting capsids (Figure 1d).

**Table 1:**
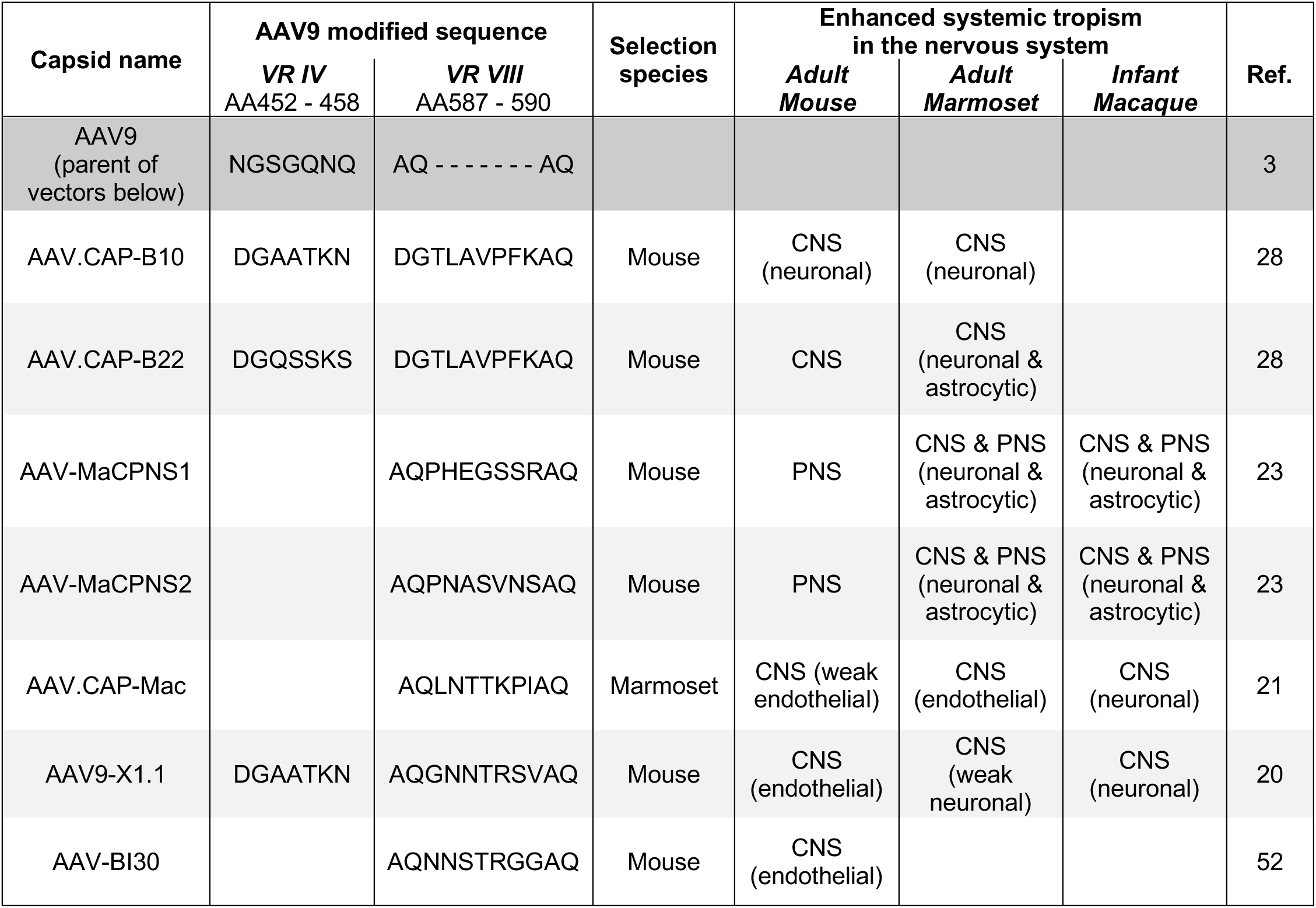
Capsid engineering details and in vivo tropisms of vectors used in this study.

After these controls, we then tested the six-capsid pool in a full screen of over 6400 proteins, including 6019 human plasma membrane proteins and secreted and cell surface-tethered proteins, as well as 397 heterodimers. We identified 22 pool hits with enhanced signal over background in each duplicate spot. To assign these hits to specific capsids in the pool, we performed follow-up deconvolution screens with each individual capsid from the pool (Figure 1e). DNA oligos for the 22 identified hits, as well as the positive control CD86, were affixed in duplicate locations to new slides. A negative control condition with no AAV analyte and a positive control condition with CTLA4-Fc (CD86 binder) were also included. We were able to successfully assign hits, including both membrane-localized and secreted proteins, to capsids. Some of these interactions were unique to specific AAV9 variants, such as LRP6 for AAV9-X1.1 or FAM234A for AAV.CAP-B22, while others were conserved across all capsids tested, such as IL3 (Table 2).

**Table 2:**
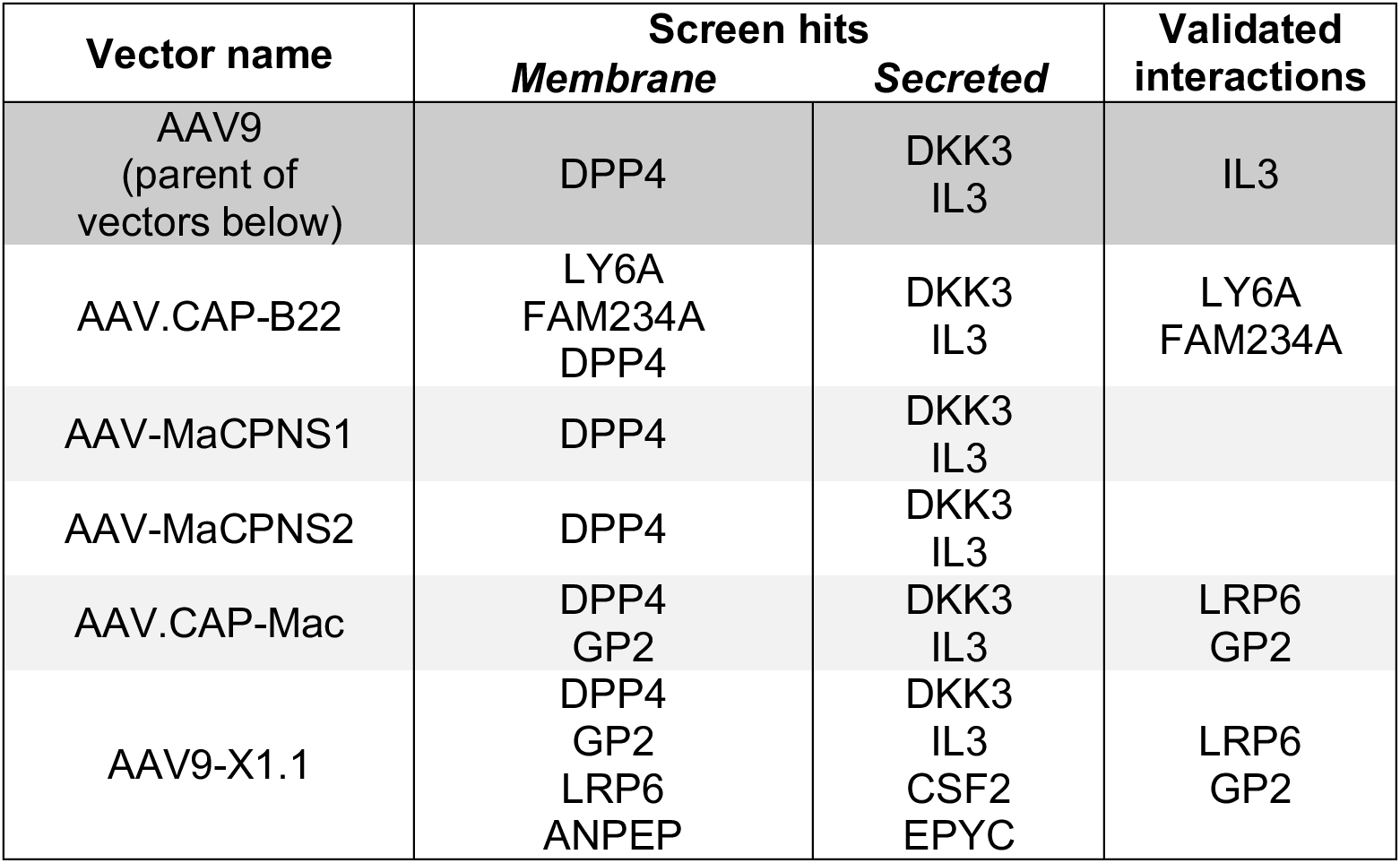
Summary of identified AAV interactions.

| Vector name | Screen hits |  | Validated interactions |
| --- | --- | --- | --- |
|  | *Membrane* | *Secreted* |  |
| AAV9<br>(parent of<br>vectors below) | DPP4 | DKK3<br>IL3 | IL3 |
| AAV.CAP-B22 | LY6A<br>FAM234A<br>DPP4 | DKK3<br>IL3 | LY6A<br>FAM234A |
| AAV-MaCPNS1 | DPP4 | DKK3<br>IL3 |  |
| AAV-MaCPNS2 | DPP4 | DKK3<br>IL3 |  |
| AAV.CAP-Mac | DPP4<br>GP2 | DKK3<br>IL3 | LRP6<br>GP2 |
| AAV9-X1.1 | DPP4<br>GP2<br>LRP6<br>ANPEP | DKK3<br>IL3<br>CSF2<br>EPYC | LRP6<br>GP2 |

### Validation of AAV binding interaction with interleukin-3

To validate binders from our cell microarray screen, hits were tested for their ability to enhance AAV potency in cell culture (Extended Data Figure 2) and capsid-binding interactions were characterized by surface plasmon resonance (SPR) (Figure 2a and Extended Data Figure 3). This reduced the candidate receptors to a subset of validated interactors (Table 2). In analyzing these interactions, we were first struck by the identified interaction of AAV9 and all its recent lab-evolved derivatives with the human immunomodulatory protein interleukin-3 (IL3) because AAVs are relatively well tolerated by the immune system^40^. IL3 is produced by activated T cells as part of the inflammatory response to viral infection, triggering expansion of various immune cells and activating type I interferon-secreting plasmacytoid dendritic cells^41^. Using SPR, we found that human IL3 binds AAV9 but not the closely related natural serotypes AAV8 and AAVrh10^42^ (Figure 2b). We then tested IL3 from different species, finding that AAV9 binds to human and macaque IL3 (83% sequence identity shared to human) but not marmoset or mouse IL3 (69% and 27% sequence identity shared with human, respectively)(Figure 2b), suggesting a binding site divergence between new and old world monkeys.

**Figure 2.**
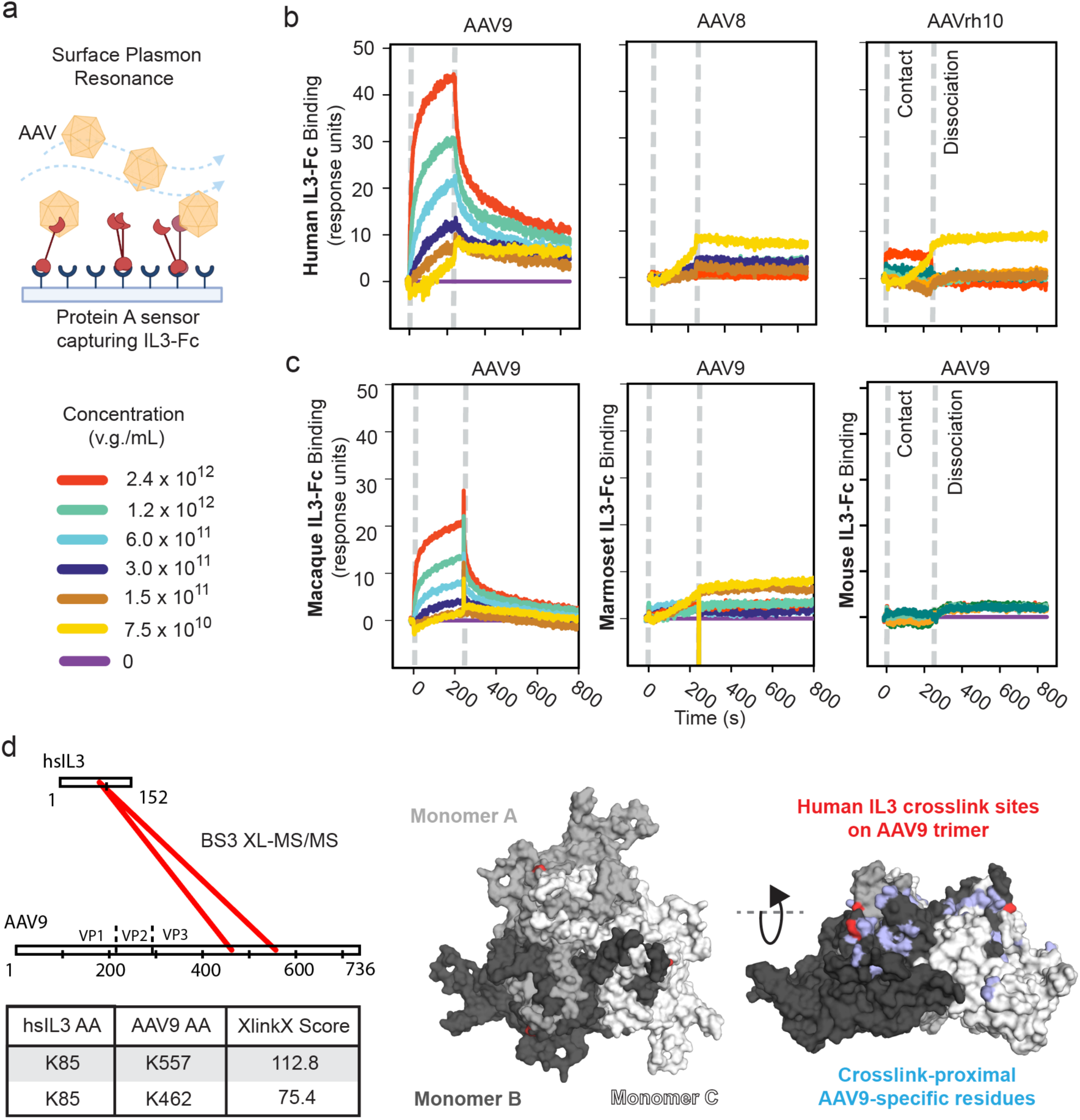
Species and serotype-specific interaction between AAV9 and the human immunomodulatory cytokine IL3. ***a***, Schematic of Surface Plasmon Resonance (SPR) experiments where IL3-Fc is captured on a protein A sensor chip and AAV analyte flows over the sensor. ***b***, SPR confirms serotype-specific interaction of AAV9 with the human immunomodulatory cytokine IL3. ***c,*** SPR confirms AAV9 binding with macaque but not marmoset or mouse IL3. ***d,*** *Left:* Scheme depicting intermolecular cross-links between human IL3 (hsIL3) and AAV9 with XlinkX scores above 40, indicating high confidence cross-link identification^76^, *Right:* Structure of AAV9 (PDB ID: 3UX1) trimer indicating human IL3 cross-linking amino acids (red) and amino acids within 20 angstroms of cross-link sites that are unique to AAV9 compared to AAV8 and AAVrh10 (blue).

**Figure 3.**
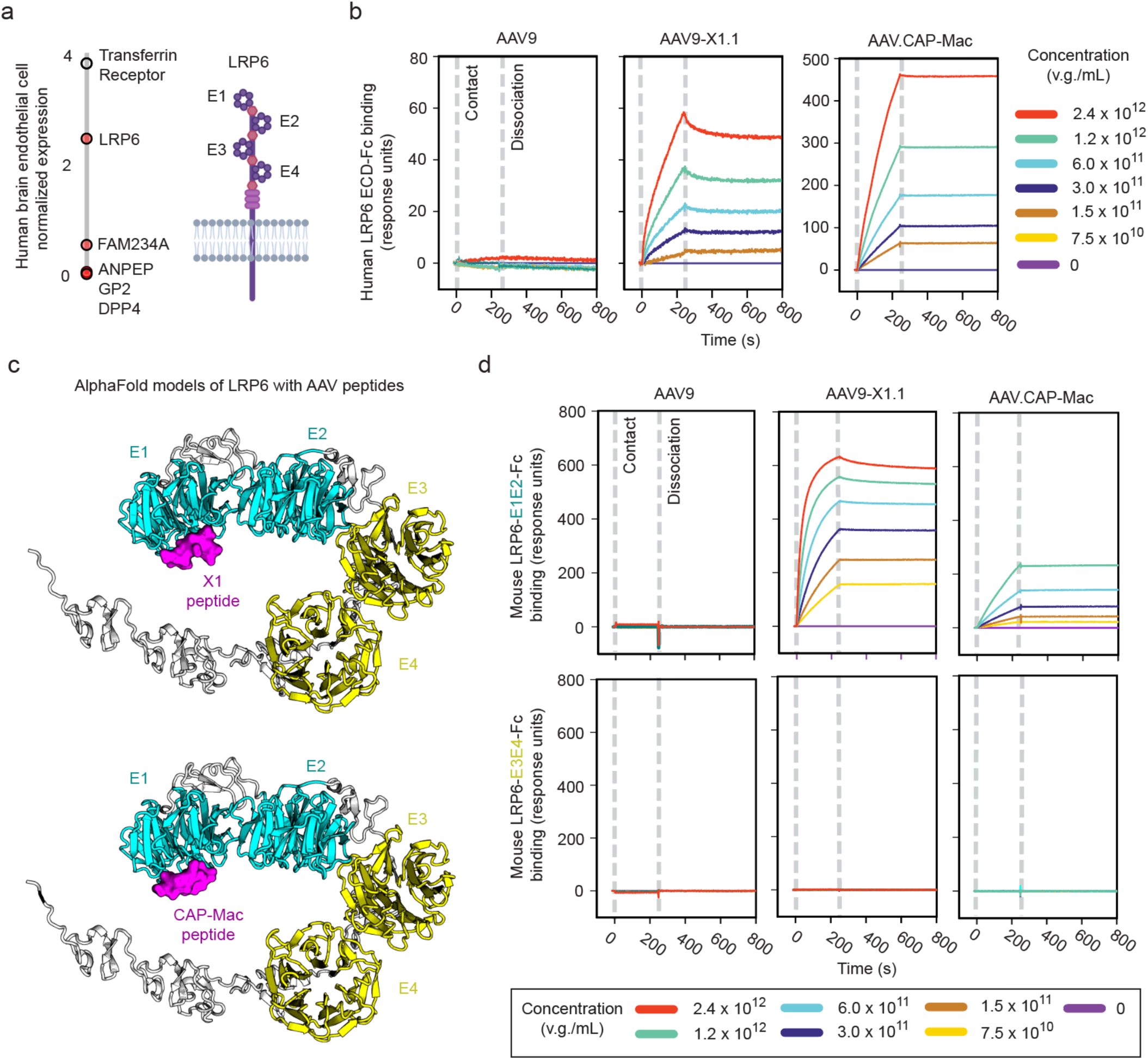
Primate brain-enhanced AAVs gain interaction with LRP6. ***a***, Arraying AAV capsid-specific hits by human brain endothelial cell expression levels reveals highly-conserved LRP6 as a potential receptor for BBB crossing. ***b***, SPR confirms that the engineered capsids AAV9-X1.1 and CAP-Mac gained direct binding interactions with human LRP6. ***c***, AlphaFold models of X1 and CAP-Mac peptides predict selective interaction with human LRP6 YWTD domain 1 (E1). ***d***, SPR of mouse LRP6-E1E2 and LRP6-E3E4 (the minimal stable extracellular domain fragments due to cooperative folding) confirms that AAV9-X1.1 and CAP-Mac bind only to LRP6-E1E2.

To further understand the species and serotype specificity of human IL3’s interaction with AAV9, we investigated the structure of the bound complex. As functional AAV ligands may have weak and dynamic monomeric interactions^43^, we leveraged avidity by flowing the 60-mer AAV9 capsid over protein A-captured dimeric IL3-Fc to ensure all biologically meaningful interactions are detected^24^. Despite this high avidity in the SPR experiment, the apparent affinity of the interaction was consistent with only a high nM interaction. Therefore, we began by performing chemical cross-linking of IL3-bound AAV9, followed by tandem mass spectrometry (XL-MS/MS)^44^. Using bis(sulfosuccinimidyl)suberate (BS3) cross-linking agent, 2 high confidence cross-links between the proteins were detected (Figure 2d, Extended Data Figure 4, and Extended Data Table 2). These cross-links place IL3’s interaction site with AAV9 near the base of the 3-fold symmetry spike and the 2-fold symmetry depression , a region containing multiple residues that are unique to AAV9 compared to non-interacting serotypes (Figure 2d).

**Figure 4.**
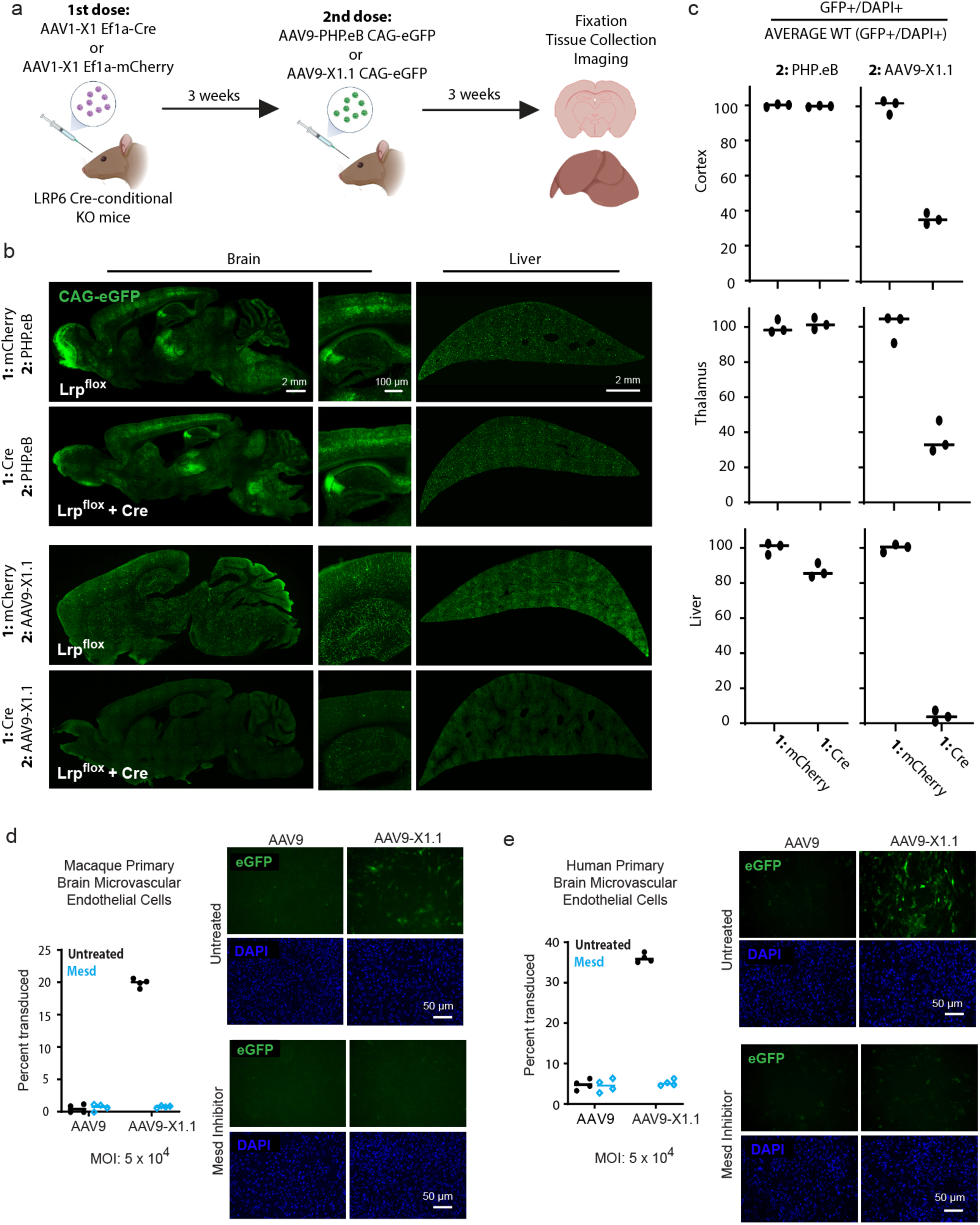
LRP6 modulates CNS function of engineered AAVs in mice. ***a***, Schematic of *Lrp6* conditional knockout by sequential AAV injection. *Lrp6* Cre-conditional knockout mice are systemically injected with AAV1-X1 packaging either Cre or mCherry, creating cohorts of mice that differ in their LRP6 expression. After allowing time for expression, these cohorts were each injected with AAV9-PHP.eB or AAV9-X1.1 packaging eGFP. By switching serotypes, neutralizing antibodies are evaded and vector dependence on *Lrp6 in vivo* may be assessed. ***b,*** Representative sagittal brain images (left) and liver images (right) from the conditional *Lrp6* knockout experiment. Imaging parameters were optimized independently for AAV9-X1.1 and AAV9-PHP.eB second dose conditions. ***c***, Quantification of AAV potency demonstrating that conditional knockout of *Lrp6* in mouse brain selectively and potently reduces AAV9-X1.1 brain and liver potency. Data points are the average of two sections per tissue region for an animal, with consistent physiological regions of interest across the four experimental cohorts. Bars represent the mean value. ***d-e***, AAV9-X1.1 has enhanced potency in ***d***, macaque and ***e,*** human primary brain microvascular endothelial cell culture, which decreases to AAV9 levels with Mesd inhibition of LRP6. Bars indicate the mean value.

### Validation of AAV binding interaction with low-density-lipoprotein-receptor-related-protein 6

Returning to the identified AAV capsid-binding interactions, we then assessed the validated AAV interactors for their potential to explain the enhanced brain tropisms of the engineered capsids. Sorting the screen hits by their expression level in endothelial cells of the human BBB^45^ spotlighted a specific interaction of low-density-lipoprotein-receptor-related-protein 6 (LRP6) with AAV9-X1.1 (Figure 3a). This capsid displays an enhanced brain endothelial-specific tropism in mice that shifts to an enhanced neuronal tropism in macaque (Table 1)^20^. Although AAV9-X1.1 contains modifications from AAV9 at both variable regions IV and VIII (Table 1), we previously showed that the tropism of AAV9-X1.1 could be transferred to other natural serotypes such as AAV1 and AAV-DJ by transferring only the variable region VIII insertion of AAV9-X1.1^20^. By SPR, here we confirm that the X1 peptide insertion in variable region VIII endows AAV1-X1 and AAVDJ-X1, but not their unmodified parent serotypes, with LRP6 binding (Extended Data Figure 5a). This demonstrates that the functional modularity of the X1 peptide in different AAV serotypes in vivo corresponds to LRP6-binding modularity.

LRP6 is a coreceptor of the canonical Wnt signaling pathway, with developmental and homeostatic roles in many tissues^46–48^. The high degree of LRP6 sequence conservation across species (98% and 99.5% sequence identity conserved between human LRP6 and mouse or macaque LRP6, respectively) aligns with AAV9-X1.1’s enhanced tropisms compared to AAV9 in rodents and primates. A similar enhancement in tropism across species is also seen for CAP-Mac^21^ (Table 1), which was engineered in marmosets and has enhanced endothelial tropism compared to AAV9 in marmosets as well as enhanced neuronal tropism in macaque. Therefore, we also tested CAP-Mac by SPR for interaction with the human LRP6 extracellular domain (Figure 3b). As with IL3, we utilized avidity to ensure that weak yet functionally important interactions are captured. Both AAV9-X1.1 and CAP-Mac strongly bind human LRP6-Fc, unlike their parent capsid, AAV9, with a sub-nM apparent affinity.

LRP6 has many endogenous WNT signaling partners with binding sites spanning either extracellular YWTD domains 1 and 2 (E1E2) or domains 3 and 4 (E3E4)^49^. We applied AlphaFold-Multimer^50^ to build models of the AAV interaction complexes, which predicted that the X1 and CAP-Mac variable region VIII peptides bind LRP6 YWTD domain 1 (Figure 3c). While the cooperative folding of E1 and E2 complicates testing of individual domains^51^, SPR results from mouse LRP6 extracellular domain fragments were consistent with the model prediction, with interaction observed for LRP6-E1E2 but not LRP6-E3E4 (Figure 3d). We also tested AAV-BI30, another engineered capsid with specific expression in the mouse brain endothelium^52^ (Table 1), finding that it also binds to LRP6-E1E2 but not LRP6-E3E4 (Extended Data Figure 5b). A pull-down assay confirmed that both AAV9-X1.1 and CAP-Mac bind the full length extracellular domain of mouse LRP6 but not that of the closely-related LRP5 (Extended Data Figure 6)^49^. AAV9, on the other hand, bound only to AAVR’s PKD2, as reported previously^39^.

AAV9-X1.1 and CAP-Mac potently infect HEK293 cells, with AAV9-X1.1 having a stronger effect (Extended Data Figure 7b). To determine if this potency is mediated by endogenous LRP6 expression in HEK293 cells, we tested the vectors with LRP6 inhibitors. AAV9-X1.1 potency was markedly reduced by Mesoderm development LRP chaperone (Mesd), a natural endoplasmic reticulum chaperone and recombinant extracellular inhibitor of LRP5 and LRP6^53^, and Sclerostin (SOST), which inhibits LRP6 through binding only E1E2^54^ (Extended Data Figure 8a and b). Importantly, neither Mesd nor SOST inhibited the potency of PHP.eB on LY6A overexpressing cells. Transient overexpression of human LRP6 boosted the potency of both capsids, with a stronger effect for CAP-Mac (Extended Data Figure 7b). This effect was largely preserved when a truncated LRP6-E1E2 was used. As expected from our pull-down assay, transient overexpression of LRP5 did not enhance the potency of either CAP-Mac or AAV9-X1.1. Together, these results support a specific functional interaction between LRP6 and both CAP-Mac and AAV9-X1.1, although the two capsids may have different functional sensitivities to LRP6 expression level. While AAV9-X1.1 more productively engages LRP6 at the lower endogenous expression levels of LRP6 in HEK293 cells, CAP-Mac shows more enhanced potency after transient overexpression of LRP6.

Of note, in addition to the intended CNS tropism receptors gained through these AAV capsid engineering efforts, both LRP6-binding engineered capsids also gained interactions with the GPI-linked protein glycoprotein 2 (GP2), which has specific pancreatic expression and, in a secreted form, plays antibacterial roles in the gut^55,56^ (Extended Data Figure 2a, Extended Data Figure 3, and Table 2). GP2 boosted the potency of both capsids in cell culture, with a stronger effect for the human protein than the mouse (Extended Data Figure 2a).

FAM234A, which bound CAP-B22 in the cell microarray screen, is found in the brain with low expression across many neuron types^57^. Although, FAM234A has been identified in disease-association studies^58^, no specific molecular function has been assigned to this protein. We found that FAM234A enhances the potency of both CAP-B22 and PHP.eB in cell culture, with the mouse receptor showing a stronger effect than the human protein (Extended Data Figure 2b). This suggests that the interaction is driven by the capsids’ shared variable region VIII insertion sequences (Table 1).

### Engineered AAVs utilize LRP6 at the blood-brain barrier in mouse and primate

Host neutralizing antibodies, developed in response to prior exposure to AAVs, complicate repeat administration with the same serotypes. Serotype-switched X1 vectors, such as AAV1-X1, were shown to enable a second systemic dosing in mice previously exposed to AAV9-based vectors^20^. We leveraged this property to determine the *in vivo* effects of AAV9-X1.1’s LRP6 interaction (Figure 4a). Brain endothelium-targeted AAV1-X1 packaging either control mCherry or Cre recombinase was systemically administered to *Lrp6* Cre-conditional knockout mice. After three weeks, either AAV9-based PHP.eB or AAV9-X1.1 packaging eGFP was systemically delivered. Whereas PHP.eB showed characteristic strong brain transduction regardless of AAV1-X1 cargo, AAV9-X1.1 brain endothelial tropism was markedly reduced in the AAV1-X1-dosed mice with *Lrp6* knocked out in AAV1-X1 transfected cells (Figure 4b and c), confirming LRP6’s necessity for capsid BBB entry *in vivo*. The AAV9-X1.1 capsid showed enhanced potency compared to PHP.eB in the liver, where LRP6 is also expressed^59^ (Extended Data Figure 8a). In *Lrp6* knockout conditions, decreased AAV9-X1.1 liver transduction was also observed (Figure 4b and c).

To confirm that LRP6 interaction is mediating the brain potency of AAV9-X1.1 in primates, we tested the vector on macaque and human primary brain microvascular endothelial cells (PBMECs) in culture (Figure 4d and 4e). AAV9-X1.1 was markedly more potent than its parent, AAV9, in the PBMECs from both species. On the other hand, the LRP6 inhibitor Mesd selectively reduced AAV9-X1.1 potency in PBMECs back to AAV9 levels. A similar LRP6-dependent boost in potency by AAV1-X1 compared to AAV1 was also observed in human PBMECs (Extended Data Figure 8b). The similarity of the responses of AAV1-X1 and AAV9-X1.1 supports our SPR experiments (Extended Data Figure 5a) that showed the X1 peptide is necessary and sufficient to target capsids to the BBB through its interactions with LRP6.

## Discussion

Recent advances in capsid engineering have led to AAV vectors that can more efficiently cross the blood-brain barrier (BBB) in rodents and non-human primates after systemic administration^17–19^, but predictable translation and further rational design of these and other non-viral BBB-crossing molecules is hampered by our limited understanding of transcytosis mechanisms, particularly in humans. This translational challenge is also an opportunity to better understand the biology of the BBB and AAV vectors. To date, only a few targets, such as transferrin receptor^60^, are used for research or therapies. Here, we developed a pipeline to find cognate receptors for engineered AAVs, focusing on the human membrane proteome and secretome. Our results validate the utility of cell microarray screening to identify receptors for natural and engineered AAVs. We identify LRP6 as a novel and highly conserved target for blood-brain barrier transcytosis by AAV9-X1.1, a potent engineered capsid for primate neurons^20^ and human IL3 as an interaction partner for AAV9. These findings offer the prospect of leveraging identified receptors for targeted drug delivery across diverse therapeutic modalities, such as small molecules, antibodies, or oligonucleotides.

Delivery vector safety and immune tolerance are key considerations with AAVs moving into the clinic as serious adverse events can occur^14–16^. Understanding the immunomodulatory potential of the IL3-AAV9 interaction we report here is therefore of high importance. Future work will have to determine whether this is a host neutralization mechanism, or a cloaking mechanism for the AAV to evade the immune system or use decoy receptors to weaken its response^61^. In addition to activated T cells, IL3 is also constitutively secreted by astrocytes in the brain to reprogram microglia, and combat Alzheimer’s disease^62^. Thus, AAV9 interaction with IL3, which is shared with all AAV9-based engineered capsids for enhanced BBB crossing, may impact processes beyond immune tolerance of the vector itself in the context of healthy and diseased brains. Importantly, the implications of this interaction cannot be readily studied in mice as AAV9, an isolate from human clinical tissue^63^, binds to human and macaque (83% AA identity) but not marmoset or mouse IL3 (69% and 27% AA identity, respectively). It is possible that this species-dependent interaction could contribute to a disconnect between rodent and primate AAV safety profiles^64–67^, especially in neurodegeneration contexts.

The high degree of sequence conservation in LRP6 (98% AA identity between mouse and human^59^) helps explain the broad conservation across species of enhanced tropism by AAV capsids targeting this receptor. We could not identify a receptor for several of the engineered capsids that we screened, such as MaCPNS1 and MaCPNS2, despite their CNS potency in rhesus macaque. It is possible that this is due to false negatives (as initially observed for CAP-Mac with LRP6), reliance on a combination of receptors only screened individually here, or these capsids utilizing a receptor whose binding site is not conserved between macaque and human. The last possibility should concern those intending to translate macaque-evolved AAVs into the clinic in the absence of mechanistic knowledge.

That both AAV9-X1.1 and CAP-Mac also showed binding to GP2, which is not present in the CNS but in the pancreas, suggests that the interaction may have piggybacked on the functional enhancement provided by LRP6-binding during directed evolution selections. This is supported by the finding that the AAVs more potently interact with human GP2 than the mouse protein that was present during the directed evolution of AAV9-X1.1. These findings highlight the importance of broad, unbiased interaction screens to build full safety profiles for engineered capsids prior to clinical trials. Surveying the diversity of mechanisms by which natural and engineered AAVs cross the BBB may also allow us to prepare defenses against future pathogens. Just as antibiotic resistance is testing our modern world, one concern is that fast-evolving pathogens will develop “BBB resistance”—the ability to access the brain and cause severe disease (as some retroviruses, including HIV-1, already do^68^). As a recent troubling example, SARS-CoV-2 capsid proteins were found in the brains of patients with long COVID, and correlated with neuropsychiatric symptoms^69^. By screening existing pathogens and their likely molecular evolutions against the growing human BBB transcytosis receptor catalog (including transferrin receptor^70,71^, insulin receptor^72,73^, CD98hc^74,75^, CA4^24^, and LRP6 (this work), we may be able to anticipate outbreaks of pathogens with neuropsychiatric sequelae.

In summary, the present study introduces a method to efficiently screen natural AAV serotypes and engineered variants against the human proteome, and expands the limited roster of targets for enhanced BBB crossing in primates. These findings suggest new strategies for successful clinical translation of engineered AAVs, provide targets for development of non-viral therapeutic modalities, and highlight latent vulnerabilities to future pathogens.

---

**Extended Data Table 1:** Doses of individual vectors within the pools used to optimize signal to noise.

| Vector name | Pool dose (v.g. / cell) |  |  |
| --- | --- | --- | --- |
| | $7.9 \times 10^5$ | $5.1 \times 10^5$ | $1.6 \times 10^5$ |
| AAV9 | $2 \times 10^5$ | $1 \times 10^5$ | $3 \times 10^4$ |
| AAV.CAP-B22 | $6 \times 10^4$ | $1 \times 10^5$ | $3 \times 10^4$ |
| AAV-MaCPNS1 | $1.5 \times 10^5$ | $1 \times 10^5$ | $3 \times 10^4$ |
| AAV-MaCPNS2 | $1.5 \times 10^5$ | $1 \times 10^5$ | $3 \times 10^4$ |
| AAV.CAP-Mac | $2 \times 10^5$ | $1 \times 10^5$ | $3 \times 10^4$ |
| AAV9-X1.1 | $3 \times 10^4$ | $1 \times 10^4$ | $1 \times 10^4$ |

**Extended Data Table 2:**
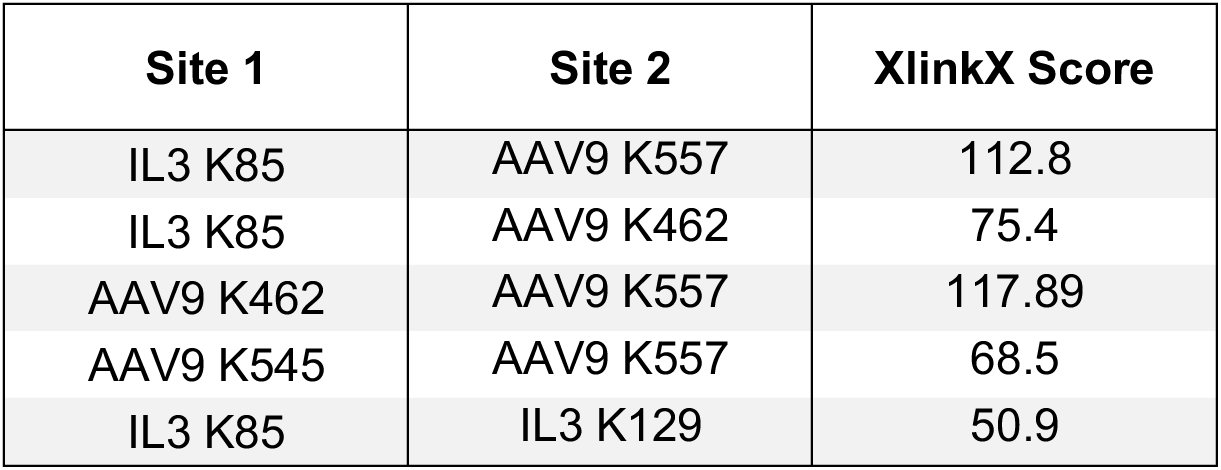
Summary of all detected BS3 cross-links with XlinkX score above 40, indicating high confidence cross-link identification in human IL3-Fc with AAV9 sample.

| Site 1 | Site 2 | XlinkX Score |
| --- | --- | --- |
| IL3 K85 | AAV9 K557 | 112.8 |
| IL3 K85 | AAV9 K462 | 75.4 |
| AAV9 K462 | AAV9 K557 | 117.89 |
| AAV9 K545 | AAV9 K557 | 68.5 |
| IL3 K85 | IL3 K129 | 50.9 |

**Extended Data Figure 1.**
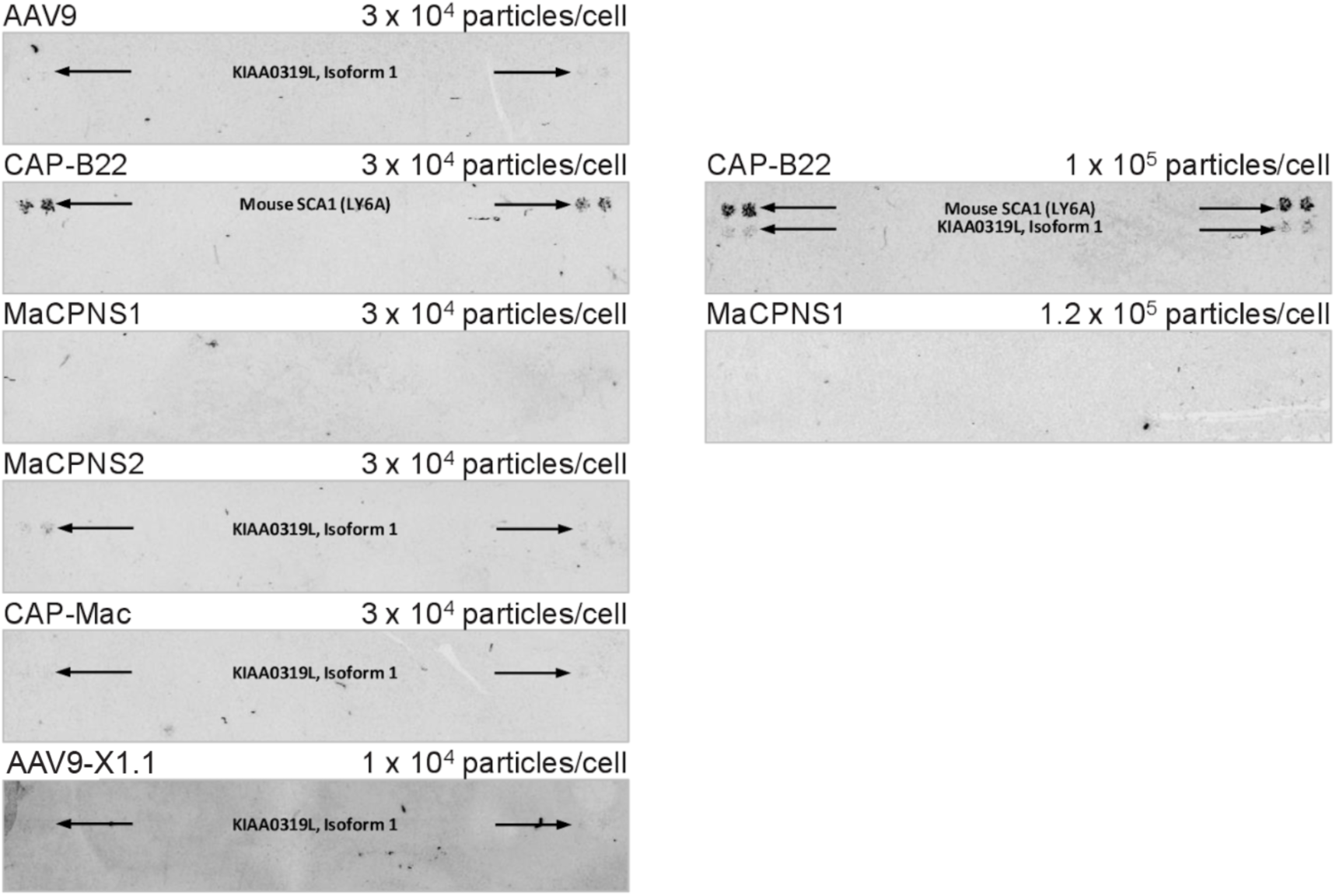
Individual characterization of pool AAVs prior to full screen. Individual AAVs were tested at various doses to determine the optimal signal to noise ratio for each capsid while confirming detection of known interactions KIAA0319L (AAVR) and, for CAP-B22 only, LY6A. AAV binding detected at duplicate spots of the same protein is indicated by arrows.

**Extended Data Figure 2.**
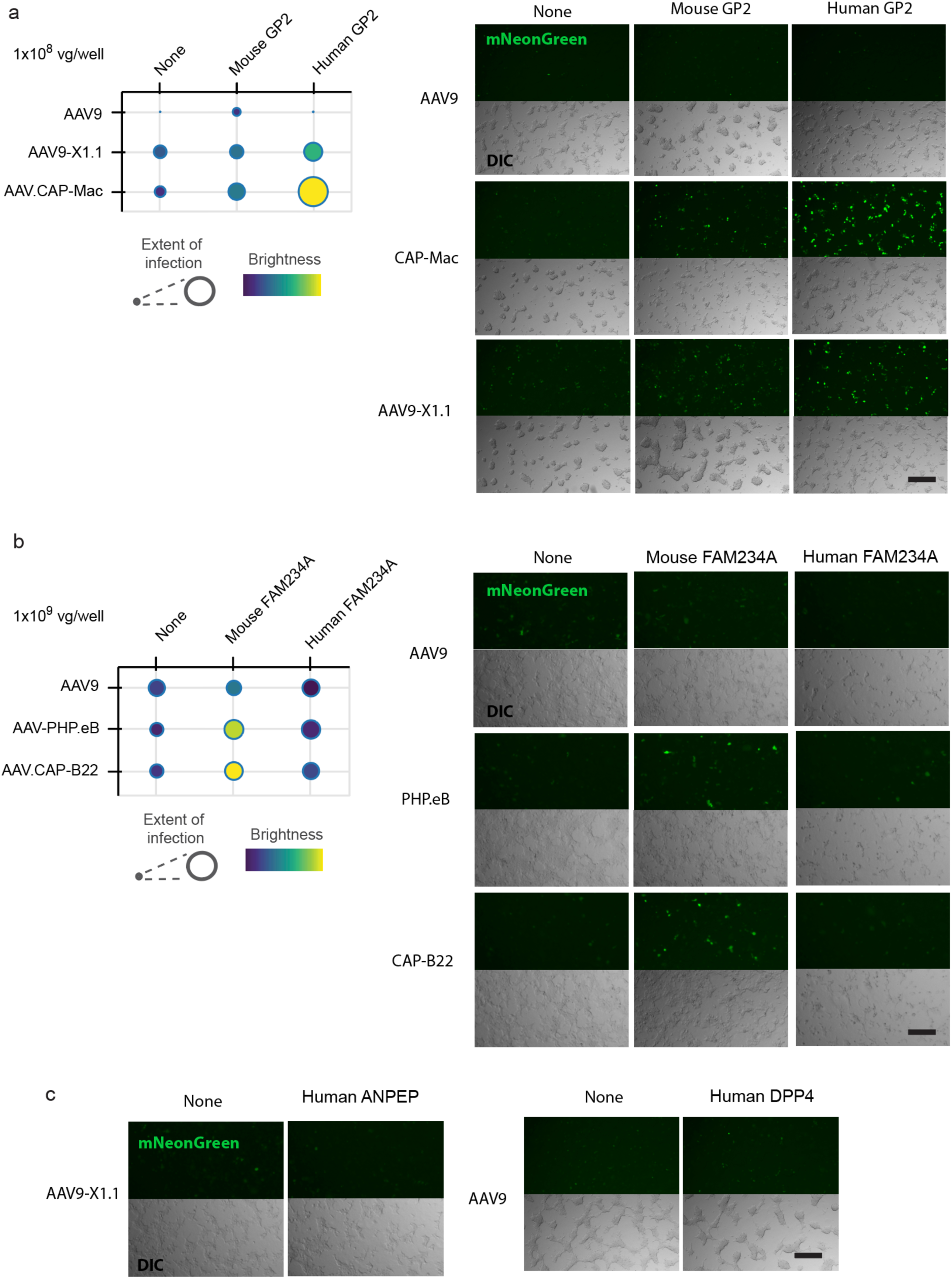
Cell culture potency assay validation of high-throughput screen hits. **a**, Transient overexpression of mouse and human GP2 in HEK293T cells resulted in enhanced potency for CAP-Mac and AAV9-X1.1, with a stronger effect for the human protein. Scales show extent of infection (min, 0.01; max, 0.17) and total brightness per signal area (min, 0.11; max, 0.45). **b**, Transient overexpression of mouse and human FAM234A in HEK293T cells results in enhanced potency for PHP.eB and CAP-B22, with a stronger effect for the mouse protein. Extent of infection (min, 0.04; max, 0.07) and total brightness per signal area (min, 0.16; max, 0.29). **c**, Transient overexpression of human ANPEP and DPP4 did not result in potency enhancements for AAV9-X1.1 and AAV9, respectively. Scale bars indicate 200 μm.

**Extended Data Figure 3.**
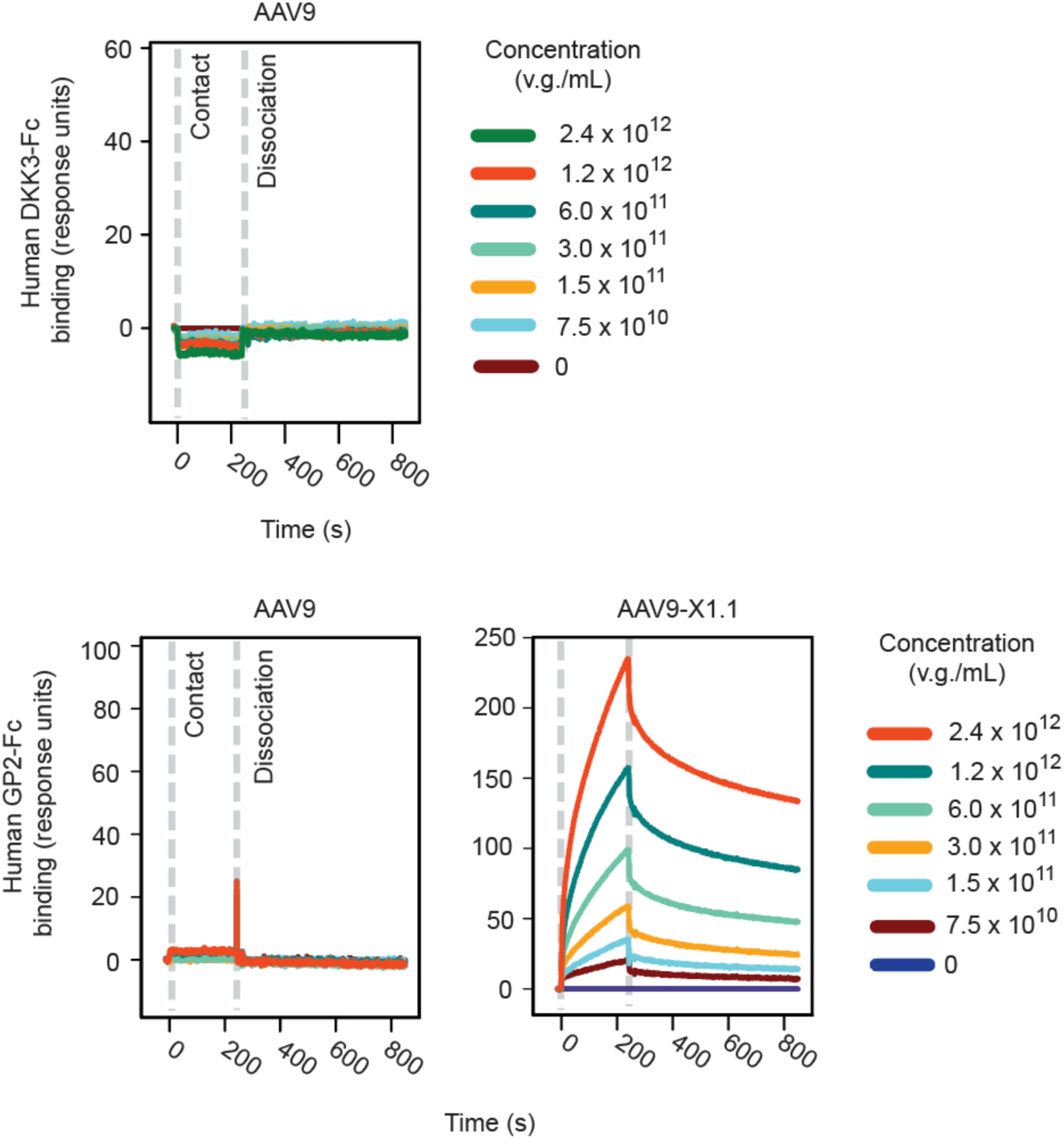
SPR confirmation of selected screen hits. Immobilization of human DKK3-Fc or human GP2-Fc on a protein A chip allowed AAV analyte interactions to be assessed. In contrast to the cell microarray screen, no interaction was observed for AAV9 with DKK3, whereas AAV9-X1.1 gained direct binding ability to human GP2, in agreement with the cell microarray screen and cell culture potency assay.

**Extended Data Figure 4.**
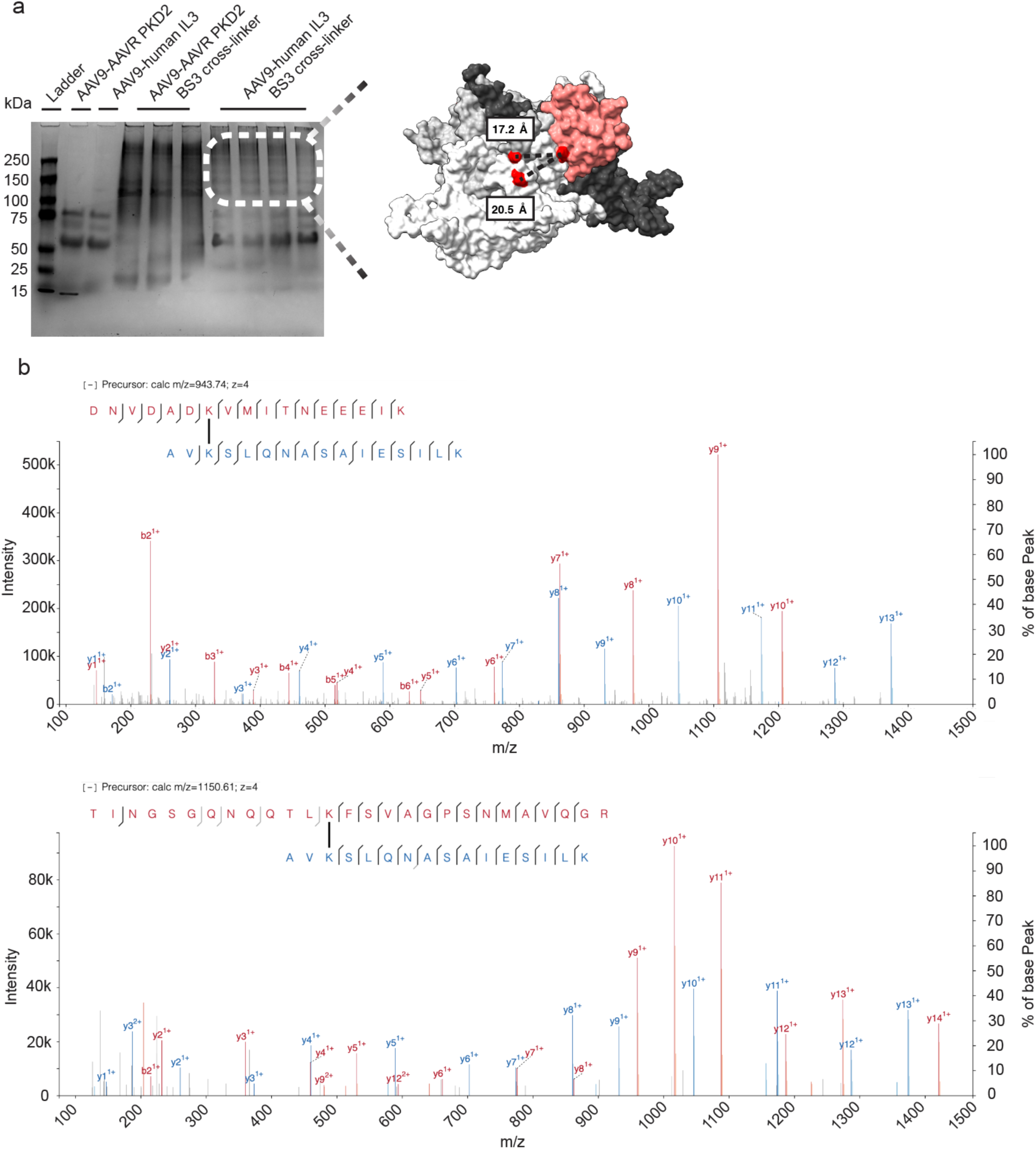
Cross-linking Mass Spectrometry confirms interaction between AAV9 and human IL3. ***a,*** PAGE gel of BS3 cross-linked AAV9 and human IL3. Dotted line indicates the region extracted for MS/MS analysis, revealing two intermolecular cross-links. These cross-links are illustrated for one potential interaction mode generated by rigid docking of human IL3 on the AAV9. AAV9 trimer in white, grey, and black. Human IL3 in red. ***b,*** Fragmentation spectra of peptides for each intermolecular cross-link between AAV9 and human IL3 with XlinkX score^76^ above 40, indicating high confidence cross-link identification. Red and blue y and b fragments indicate the peptide of origin presented in the upper left.

**Extended Data Figure 5.**
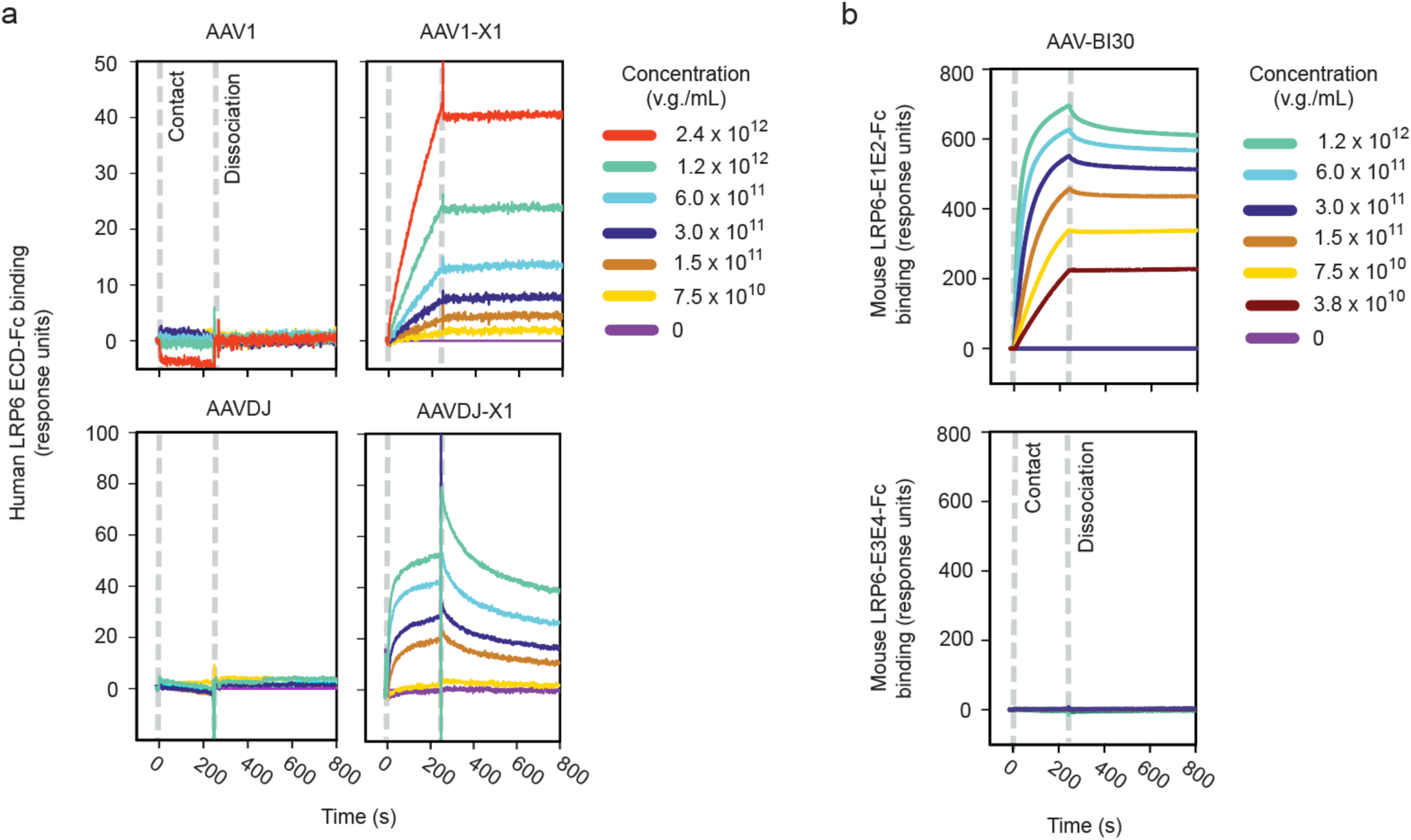
LRP6 binding to X1 peptide in multiple serotypes and to AAV-BI30. **a**, SPR of the complete human LRP6 extracellular domain confirmed that the X1 insertion peptide modularly enables LRP6 binding across multiple serotypes. **b,** SPR of AAV-BI30 confirmed binding interaction with mouse LRP6-E1E2 and not LRP6-E3E4.

**Extended Data Figure 6.**
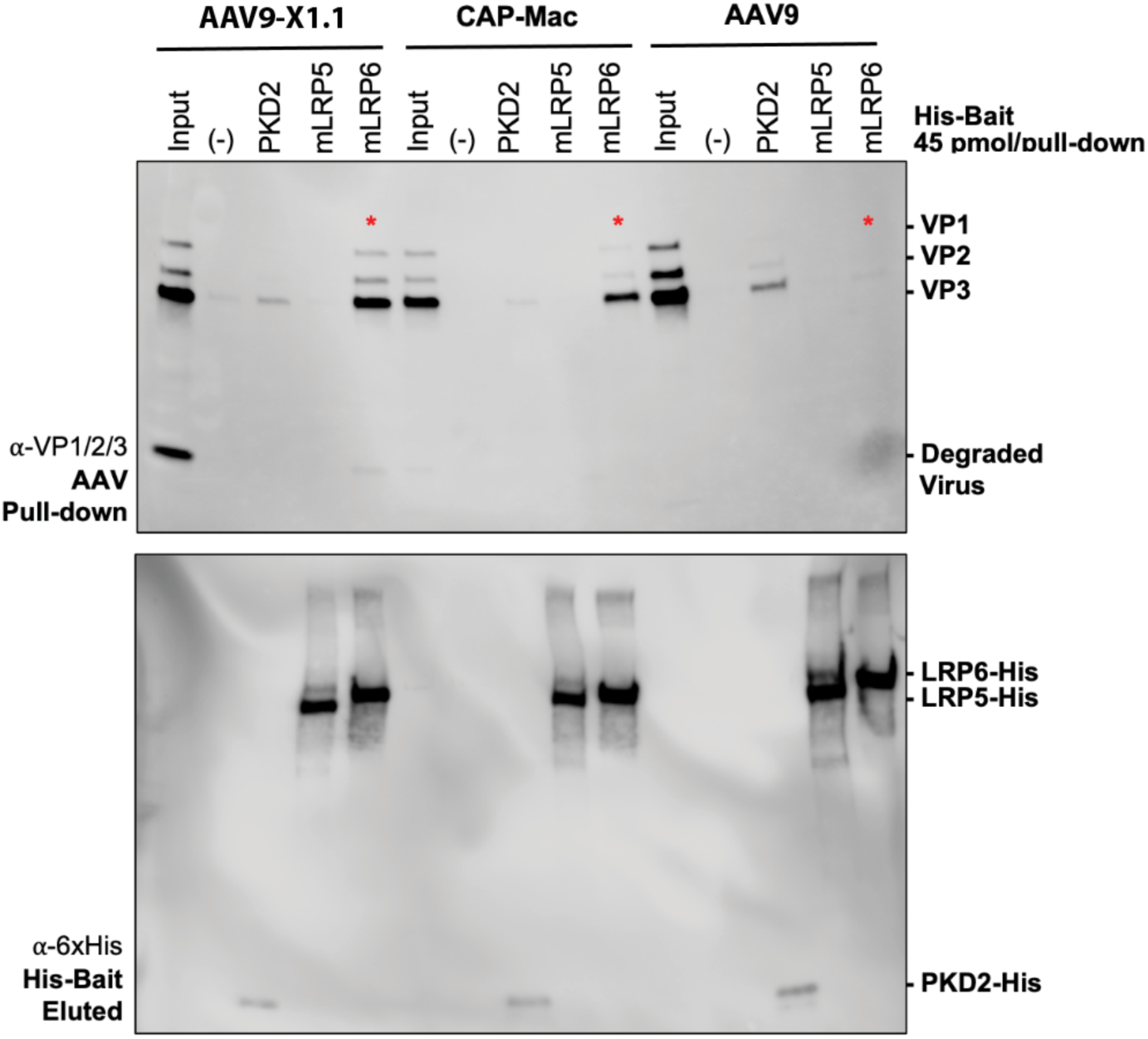
AAV9-X1.1 and CAP-Mac bind LRP6 and AAVR but not LRP5. Pull-down assay with the extracellular domains of mouse LRP6, mouse LRP5, and human AAVR PDK2 domain against AAV9, AAV9-X1.1, and CAP-Mac prey. Red asterisks indicate the LRP6 binding interaction gained by X1.1 and CAP-Mac during directed evolution from parent capsid AAV9.

**Extended Data Figure 7.**
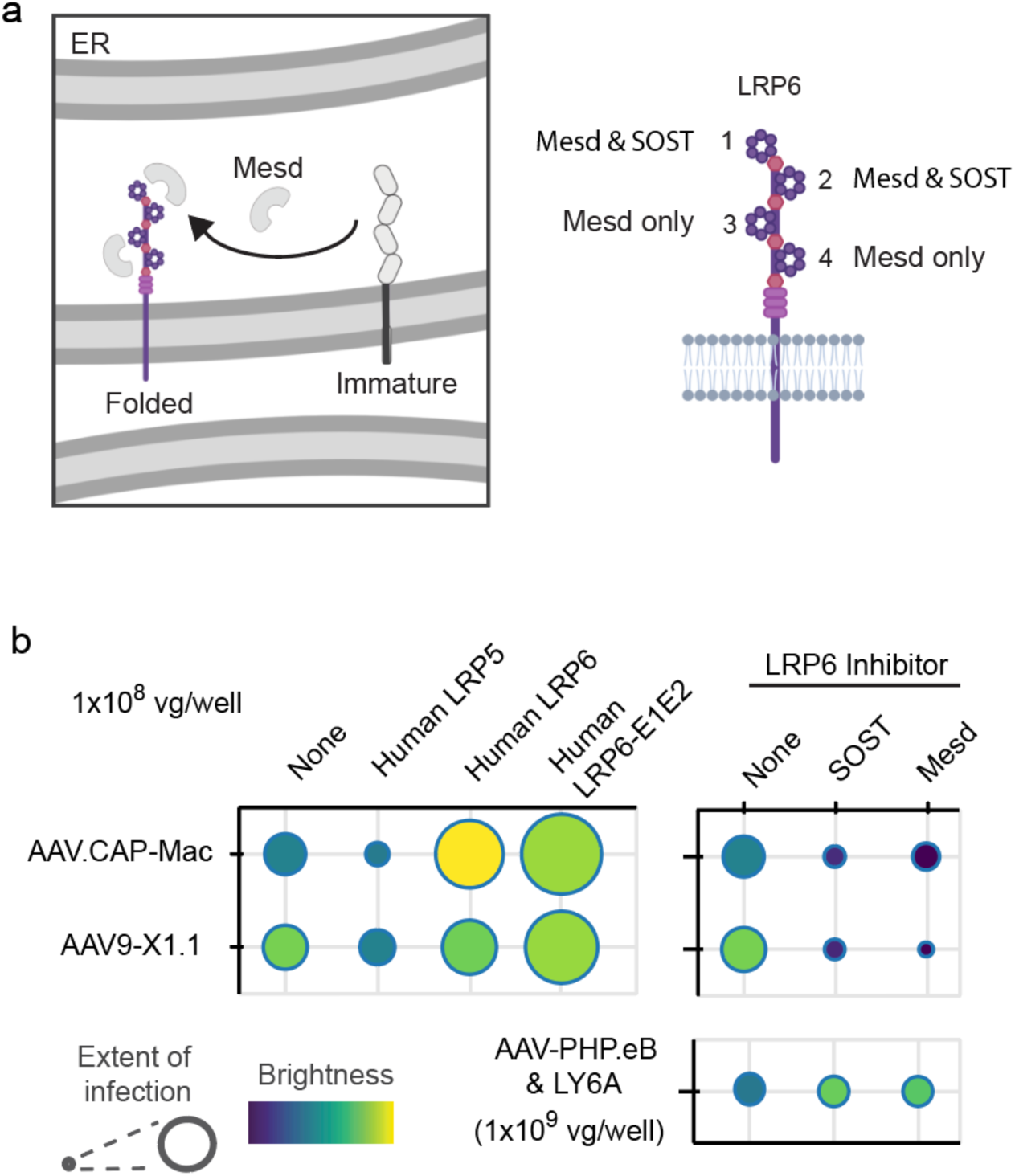
Cell culture potency assay validation of LRP6 interaction. **a**, Schematic of Mesd chaperone function and LRP6 domain-dependent inhibition by recombinant Mesd and SOST proteins. **b,** Quantification of AAV potency demonstrating the effects of LRP receptor transient overexpression and LRP6 inhibition. Extent of infection (min, 0.04; max, 0.23) and total brightness per signal area (min, 0.04; max, 0.51)

**Extended Data Figure 8.**
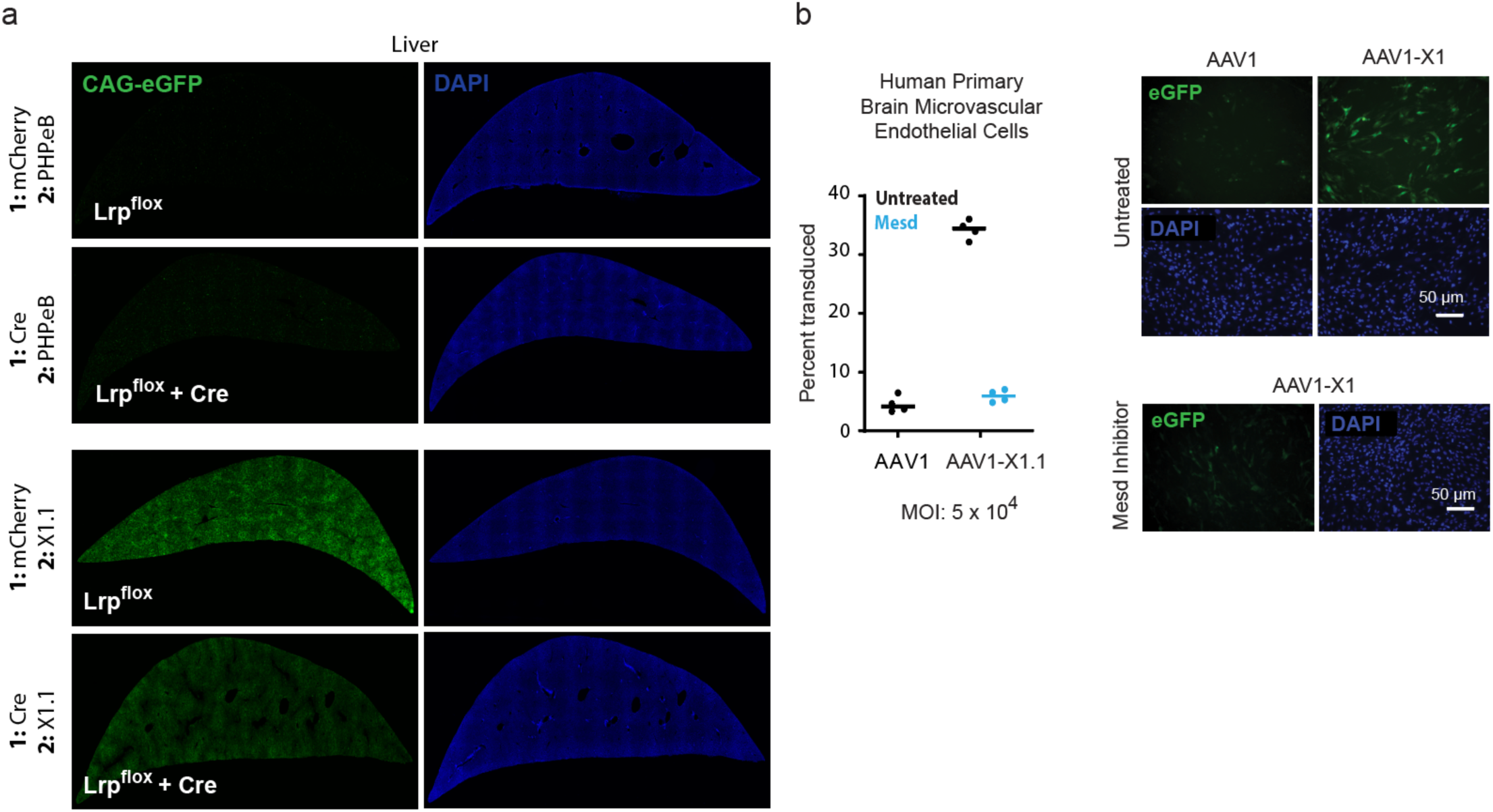
Potency of X1 AAVs in mouse liver and human primary cell culture. **a**, AAV9-X1.1 has stronger potency in mouse liver than PHP.eB. Representative liver images from Figure 4b with imaging parameters re-optimized for AAV9-X1.1 LRP^flox^ and applied to all conditions. **b,** AAV1-X1 has enhanced potency in human primary brain microvascular endothelial cell culture, which decreases to AAV1 levels with Mesd inhibition of LRP6. Bars indicate the mean value.

## Methods

### Viral vector production

AAVs were produced as previously described^77^. Briefly, HEK293 cells were triple transfected with capsid, genome, and helper plasmids. Media was exchanged the next day then collected and replaced two days after. At five days post-transfection, media and cells were collected and processed for AAV purification. Cells were lysed in a high salt solution and treated with salt-activated nuclease. Media were PEG precipitated and resuspended in salt-activated nuclease solution. Both solutions were added to iodixanol density columns, ultracentrifuged, and extracted from the 40%/60% interface. Finally, AAVs were buffer exchanged, concentrated, titered, and (for vectors destined for non-human primates) assayed for endotoxin using Piece LAL chromogenic endotoxin kit (cat# A39552).

### Retrogenix cell microarray

Retrogenix cell microarray was performed as previously described^31,32^ with the following adaptations for AAV analytes. Pre-screen optimizations were performed on slides of HEK293 cells and cells overexpressing mouse LY6A and human AAVR (KIAA0319L), TGFBR2, and EGFR. Transfection efficiencies were validated to exceed a minimum threshold prior to analyte application. AAVs were added to fixed cells at a concentration of 6 x 10^4^ AAV particles per HEK293 cell.

Biotinylated AAVs were created by incubation for 2 hours at room temperature with 10,000-fold molar ratio of NHS-PEG4-biotin (Thermo A39259) to AAV at 1x10^13^ viral genomes (v.g.) per mL in PBS. Reactions were quenched with 1 M Tris, pH 8 prior to buffer exchange, concentration, and AAV re-titer. Biotinylated AAVs were detected on HEK293 cells post fixation by AF647-labeled streptavidin. Unlabeled AAVs were detected on HEK293 cells post fixation by anti-AAV9 clone HL2372 (Merck, MABF2309-100UL) at a 1:500 dilution followed by AF647-labeled anti mIgG H+L.

To achieve a suitable signal to noise ratio, necessary for minimizing false positives and false negatives, unlabeled AAVs were screened individually and as a pool at various concentrations, using anti-AAV9 detection. The final test pool was screened against fixed HEK293 cells/slides expressing approximately 6000 human plasma membrane proteins, secreted and cell surface tethered human secreted proteins and approximately 400 human heterodimers, each in duplicate. Hits were identified using ImageQuant as spots observed in duplicate. Following the screen, the 22 identified hits and CD86 positive control protein were spotted on new slides for individual AAV testing in a deconvolution screen. A negative control condition with no analyte and a positive control condition with CTLA4-Fc (to interact with CD86) were also included.

### Protein preparation

Lyophilized mouse LRP6 (AA20-1366) tagged with 6xHis tag, N-terminal (E1E2) and C-terminal half (E3E4) fragments of mouse LRP6 (N-half: AA 20-628, C-half: AA 629-1244) tagged with Fc (mouse IgG2a), and full-length human LRP6 (AA 20-1368) tagged with Fc (human IgG1), and LRP5 (AA1-1383) tagged with 6xHis tag, and SOST protein were purchased from Bio-Techne (cat# 2960-LR-025, 9950-LR-050, 9954-LR-050, 1505-LR-025, 7344-LR-025/CF, 1406-ST, respectively). Mesd protein was purchased from SinoBiological (cat# 10949-H08H). All proteins were reconstituted in Dulbecco’s phosphate buffered saline (DPBS, Gibco^TM^) at desired concentrations before use.

Human, macaque, marmoset, and mouse Interleukin 3 (hIL3: AA 1-152, macaque IL3: AA 1-144, marmoset IL3: AA 1-143, mIL3: AA 1-166) triple tagged with Fc-Myc-6xHis, human and mouse GP2 (hGP2: AA 1-518, mGP2: AA 1-515) triple tagged with Fc-Myc-6xHis, and human and mouse DKK3 (hDKK3: AA 1-350, mDKK3: AA 1-349) triple tagged with Fc-Myc-6xHis were transfected into Expi293F^TM^ (Thermo Fisher Scientific) cells at a density of 3 × 10^6^ viable cells/mL using ExpiFectamine^TM^ (Thermo Fisher Scientific) according to the manufacturer’s manual, and secreted proteins in media were harvested after 120 hours and cleared using a 0.45-µm PVDF vacuum filter (Sigma Millipore). Each His-tagged protein in media was captured with Ni-NTA resin (Qiagen) and eluted with DPBS containing 150 mM imidazole.

Human Adeno-Associated Virus Receptor (AAVR) PKD2 domain (AA 401-498) tagged with 6xHis was purified as described previously^39^. Briefly, PKD2 was expressed in BL21(DE3)-RIPL *E. coli*. Cells were lysed by sonication, and the insoluble fraction was cleared by centrifugation. Cleared lysate was applied to a Ni-NTA column (Qiagen) and eluted using DPBS containing 250 mM imidazole.

### Surface Plasmon Resonance (SPR)

A Sierra SPR-32 (Bruker) loaded with a protein A sensor chip was used. Fc-fusion proteins in HBS-EP+ buffer (GE Healthcare) were immobilized at a capture level of 600-800 response units (RU) for Figure 2b, 2c, 3d, Extended Data Figure 3, and Extended Data Figure 5, and 1200-1500 RU for Figure 3b. AAVs were injected at a flow rate of 10 µL per min for 240 seconds followed by a 600 second dissociation. AAV concentrations began at 2.4 x 10^12^ v.g. per mL and proceeded at 2-fold dilution intervals. A regeneration step with 10 mM glycine pH 1.5 was performed between each cycle. All kinetic data were double reference subtracted.

### Pull-down assay

The pull-down assay was performed as described previously^39^. Briefly, prey AAVs were mixed with His-tagged bait protein and Ni-NTA resin in a binding buffer of DPBS containing 20 mM imidazole for 1 h at 4°C on an orbital mixer. Resin was then collected in a spin column, washed twice with 10 column volumes of binding buffer and eluted in 45 μL of DPBS containing 150 mM imidazole. Eluate was analyzed by Western blot using anti-VP1/VP2/VP3 (ARP, cat# 03-61058) and anti-6xHis (Abcam, cat# ab18184) antibodies.

### HEK293 cell culture potency assay

HEK293T cells in Dulbecco’s Modified Eagle Medium (DMEM) containing 5% fetal bovine serum (FBS), 1% non-essential amino acids (NEAA), and 100 U per mL penicillin-streptomycin were cultured in 6-well plates at 37°C in 5% CO2. At 80% confluency, cells were transiently transfected with 2.53 µg plasmid DNA encoding a membrane protein hit from the Retrogenix cell microarray screen. Cells were transferred to 96-well plates at 20% confluency and maintained in FluoroBrite^TM^ DMEM supplemented with 0.5% FBS, 1% NEAA, 100 U per mL penicillin-streptomycin, 1x GlutaMAX, and 15 µM HEPES. Plates were imaged 24 hours after application of AAV on a Keyence BZ-X700 (4x objective). For experiments with protein inhibitors, Mesd (26 ug/ml) and SOST (0.2 µg/ml) were added 4 hours prior to AAV addition. NucBlue^TM^ Live ReadyProbes^TM^ reagent (Hoechst 33342) was added to each well to aid autofocusing. Image quantification was performed as described previously^24^, using our custom Python image processing pipeline, available at github.com/GradinaruLab/in-vitro-transduction-assay.

### Cross-linking Mass Spectrometry

The cross-linking procedure was modified from the manufacturer’s instructions (Thermo Fisher). In brief, purified AAV9 was complexed with purified hIL3 at a 1:2 ratio of 300 μM AAV & 600 μM hIL3, respectively. Bis(sulfosuccinimidyl)suberate (BS3) (Thermo Fisher) was added to a final concentration of 3 mM and incubated at room temperature for 1 hour. After incubation, Tris buffer was added to a final concentration of 20 mM to quench the reaction. The samples were then run on an SDS poly-acrylamide gel and stained with Coomassie blue. Bands corresponding to cross-linked protein were cut out of the gel and further processed for mass spectrometry analysis.

The cut-out gel bands were washed with 50 mM NH4HCO3 in 50% acetonitrile and dehydrated with 100% acetonitrile before drying. The dried sample was reduced with 10 mM DTT and then alkylated with 100 mM chloroacetamide. The sample was then dehydrated with acetonitrile and dried before overnight digestion with a 20 ng per μL solution of trypsin in 50 mM NH4HCO3. Digestion was arrested with 5 μL of 5% formic acid. The sample was then centrifuged and the supernatant containing digested peptides was collected. Digested peptides were then desalted using ZipTip according to the manufacturers protocol (Millipore). Desalted peptides were then eluted, dried, and then suspended in LC-MS-grade water containing 0.2% formic acid and 2% acetonitrile for LC-MS/MS analysis. LC-MS/MS analysis was performed with an EASY-nLC 1200 (ThermoFisher Scientific, San Jose, CA) coupled to a Q Exactive HF hybrid quadrupole-Orbitrap mass spectrometer (ThermoFisher Scientific, San Jose, CA). Peptides were separated on an Aurora UHPLC Column (25 cm × 75 μm, 1.7 μm C18, AUR3-25075C18, Ion Opticks) with a flow rate of 0.35 μL/min for a total duration of 43 min and ionized at 2.2 kV in the positive ion mode. The gradient was composed of 6% solvent B (2 min), 6-25% B (20.5 min), 25-40% B (7.5 min), and 40–98% B (13 min); solvent A: 2% acetonitrile and 0.2% formic acid in water; solvent B: 80% acetonitrile and 0.2% formic acid. MS1 scans were acquired at the resolution of 60,000 from 375 to 1500 m/z, AGC target 3e6, and maximum injection time 15 ms. The 12 most abundant ions in MS2 scans were acquired at a resolution of 30,000, AGC target 1e5, maximum injection time 60 ms, and normalized collision energy of 28. Dynamic exclusion was set to 30 s and ions with charge +1, +7, +8 and >+8 were excluded. The temperature of ion transfer tube was 275°C and the S-lens RF level was set to 60.

For cross-link identification, MS2 fragmentation spectra were searched and analyzed using Sequest and XlinkX nodes bundled into Proteome Discoverer (version 2.5, Thermo Scientific) against *in silico* tryptic digested protein sequences including AAV9 capsid protein VP1 and Interleukin-3 retrieved from Uni-Prot (Q6JC40 and Q6NZ78, respectively). The maximum missed cleavages was set to 2. The maximum parental mass error was set to 10 ppm, and the MS2 mass tolerance was set to 0.05 Da. For BS3 cross-links, variable cross-link modifications were set as DSS (K and protein N-terminus, +138.068 Da) and the dynamic modifications were set as DSS hydrolyzed on lysine (K, +156.079 Da), oxidation on methionine (M, +15.995 Da), protein N-terminal Met-loss (-131.040 Da) and protein N-terminal acetylation (+42.011 Da). Carbamidomethylation on cysteine (C, +57.021 Da) was set as a fixed modification. The false discovery rate (FDR) for cross-linked peptide validation was set to 0.01 using the XlinkX/PD Validator Node and cross-links with Xlinkx score^76^ greater than 40 were reported here. Mass spectrometry proteomics data have been deposited to the ProteomeXchange Consortium via the PRIDE^78^ partner repository with the dataset identifier PXD045380. Identified cross-links were visualized using xiSPEC^79^.

### Primary cell culture potency assay

Human brain microvascular endothelial cells (ScienCell Research Laboratories, cat# 1000) and cynomolgus monkey primary brain microvascular endothelial cells (CellBiologics, cat# MK-6023) were cultured as per the instructions provided by the vendor. The cell cultures were then treated with single-stranded AAV genome CAG-eGFP packaged viral vectors at a multiplicity of infection (MOI) of 5x10^4^ per well (4 wells per vector). The fluorescence expression of the culture was inspected and quantified one day after the infection procedure.

### Animals

All mouse procedures were approved by the California Institute of Technology Institutional Animal Care and Use Committee (IACUC). Adult (6-8 weeks old) homozygous B6;129S-Lrp6tm1.1Vari/J mice (Jackson Labs #026267) were retro-orbitally administered with 1 x 10^12^ v.g. per animal of AAV1-X1 packaging either Ef1a-mCherry or Ef1A-Cre (N=6 per condition). After 3 weeks, those mice were re-administered with 1 x 10^12^ v.g. per animal of PHP.eB or AAV-X1.1 packaging CAG-eGFP (N=3 per condition). Mice were randomly assigned to a particular AAV condition. Experimenters were not blinded for any of the experiments performed in this study.

### *Lrp6* conditional knockout tissue preparation and imaging

Mice were anesthetized with Euthasol (pentobarbital sodium and phenytoin sodium solution, Virbac AH) and transcardially perfused with about 50 mL of 0.1 M PBS, pH 7.4 followed by an equal volume of 4% paraformaldehyde (PFA) in 0.1 M PBS. Collected organs were post-fixed in 4% PFA overnight at 4°C, washed, and stored in 0.1 M PBS with 0.05% sodium azide at 4°C. A Leica VT1200 vibratome was used to prepare 100 um brain sections to be imaged on a Zeiss LSM 880 confocal microscope using a Plan-Apochromat 10x 0.45 M27 (working distance, 2.0 mm) objective. Images were analyzed in Zen Black 2.3 SP1 (Zeiss) and ImageJ.

### AlphaFold structure modeling

The complex structures of LRP6 ECD and AAV-X1 or AAV.CAP-Mac VR-VIII peptide were modeled using a cloud-based implementation of AlphaFold-Multimer-v3^50^ provided in ColabFold v2.3.5^80^. The input comprised two sequences: surface-exposed residues in VR-VIII of AAV-X1 (587-AQGNNTRSVAQAQTG-594) or AAV-CAP-Mac (587-AQLNTTKPIAQAQTG-594) and the extracellular domain of human LRP6 (UniProt entry O75581, residues 20-1370). We ran the Google Colaboratory notebook using an A100 SXM4 40GB GPU. Five structure models were produced using a protocol with up to 20 recycles, and MSA generated with MMseqs2 (UniRef+Environmental)^81^ and templates from PDB70. The structure models were ranked using a weighted combination of pTM and iPTM scores as described in^50^.

## Acknowledgements

We thank Catherine Oikonomou for help with manuscript editing. We thank Helen McBride for assistance in establishing collaborations with Charles River Laboratories, Brad Gartland and Lyndsey Chatham of Charles River Laboratories for technical assistance, and Máté Borsos for assistance breeding *Lrp6* conditional KO mice and providing AAV8 and AAVrh10. We thank Nathan Appling for helpful discussion. Cryo-electron microscopy was perfomed in the Beckman Institute Resource Center for Transmission Electron Microscopy at Caltech.

Cross-linking mass spectrometry was performed in the Beckman Institute Proteome Exploration Laboratory. Figures were created using imagery from Biorender. This project was supported by the Beckman Institute CLOVER Center (to T.F.S. and V.G.) and NIH PIONEER DP1NS111369 (to V.G.), and NIH BRAIN Initiative Armamentarium UF1MH128336 (to V.G. and T.F.S.).

## Author Contributions

T.F.S. and V.G. conceived the project. T.F.S., X.C., and V.G. wrote the manuscript and prepared figures with input from all authors. T.F.S., X.C., E.E.S., Y.L., S.J., and M.R.C. produced AAVs. B.W. and C.T. performed cell microarray screening. S.J. produced recombinant receptor protein and performed pull-down assays, T.J.B., T.Y.W., and T.F.C. performed cross-linking mass spectrometry experiments. C.M.A. and E.E.S performed cell culture potency assays. T.F.S. and S.J. performed SPR experiments. X.D. performed AlphaFold modeling. X.C. and D.A.W. performed mouse experiments. X.C. performed primary cell culture experiments. T.F.S and V.G. supervised and V.G. funded the project.

## Competing Interests

The California Institute of Technology has filed a provisional patent on this work with T.F.S, X.C., S.J., and V.G. listed as inventors. V.G. is a co-founder and board of directors member of Capsida Therapeutics, a fully integrated AAV engineering and gene therapy company. B.W. and C.T. are employees of Charles River Laboratories. The remaining authors declare no competing interests.

